# Changing word meanings in biomedical literature reveal pandemics and new technologies

**DOI:** 10.1101/2022.06.27.497742

**Authors:** David N. Nicholson, Faisal Alquaddoomi, Vincent Rubinetti, Casey S. Greene

## Abstract

While we often think of words as having a fixed meaning that we use to describe a changing world, words are also dynamic and changing. Scientific research can also be remarkably fast-moving, with new concepts or approaches rapidly gaining mind share. We examined scientific writing, both preprint and pre-publication peer-reviewed text, to identify terms that have changed and examine their use. One particular challenge that we faced was that the shift from closed to open access publishing meant that the size of available corpora changed by over an order of magnitude in the last two decades. We developed an approach to evaluate semantic shift by accounting for both intra- and inter-year variability using multiple integrated models. Using this strategy and examining year-by-year changes revealed thousands of change points in both corpora. We found change points for tokens including ‘cas9’, ‘pandemic’, and ‘sars’ among many others. The consistent change-points between pre-publication peer-reviewed and preprinted text were largely related to the COVID-19 pandemic. We developed a web app for exploration (https://greenelab.github.io/word-lapse/) that enables users to investigate individual terms. To our knowledge, this analysis is the first to examine semantic shift in biomedical preprints and pre-publication peer-reviewed text, and it lays the foundation for future work to examine how terms acquire new meaning and the extent to which that process is encouraged or discouraged by peer review.

## Introduction

Language is constantly evolving, and the meaning that we ascribe to words changes over time. For example, the word “nice” was used to mean foolish or innocent back in the 15th-17th century; then, it underwent a positive shift to its current meaning of “pleasant or delightful”[1]. These shifts occur for many reasons. For example, writers may use new metaphors or substitute words for others with similar meanings in a process known as metonymy [1]. Studying these shifts can provide a nuanced understanding of how language adapts to describe our world.

Scientific fields of inquiry also change, sometimes rapidly, as researchers devise and test new hypotheses and applications. For example, the repurposing of the CRISPR-Cas9 system to a pervasive tool for genome editing has altered how we discuss molecular entities. Microbes use this as an immune system to defend against viruses. Scientists repurposed this system for genome editing [2], leading to changes in the use of the term. Science is a field with substantial written communication [3], both via published papers [4] and preprints [5,6]. Examining scientific manuscripts with computational linguistics can reveal longitudinal trends in scientific research.

Studying changes in the use of word meanings is called semantic shift detection. Approaches for semantic shift detection examine time series datasets that capture word usage patterns, both with respect to frequency and structure. Typically, these time series are generated for individual words by training a unique model on text binned by a selected time period [7,8,9]. Methods are then applied to identify “change points” where a word’s meaning has changed [11].

Semantic shifts have been examined in many sources. Analysis has included newspapers [12,13,14], books [7], reddit [15], and Twitter [16]. Researchers have examined topics in information retrieval [17], and in biomedicine COVID-19 has been examined multiple times [18,19,20]. The amount of open access biomedical literature has dramatically increased in the last two decades, laying the groundwork for the large-scale analysis of semantic shifts in biomedicine.

We examine these semantic shifts in this rapidly growing body of open access text. We include both published papers and preprints in our analysis. We found that novel strategies integrating multiple models for each year sidestepped the challenge of instability in the machine learning models and allowed us to estimate intra- and inter-year variability. We identify semantic change points for each token. We examine key cases and provide the full set of research products, including change points and machine learning models, as openly licensed tools for the community. We also created a webserver that allows users to analyze tokens of interest on the fly, examining both the most similar terms within a year and temporal trends.

## Methods

### Biomedical Corpora Examined

#### Pubtator Central

Pubtator Central is an open-access resource containing annotated abstracts and full-text annotated with entity recognition systems for biomedical concepts [21]. The methods used are TaggerOne [22] to tag diseases, chemicals, and cell line entities, GNormPlus [23] to tag genes, SR4GN [24] to tag species, and tmVar [25] to tag genetic mutations. We initially downloaded this resource on December 07th, 2021, and processed over 30 million documents. This resource contains documents that date back to the pre-1800s to the year 2021; however, due to the low sample size in early years, we only used documents published from 2000 to 2021. The resource was subsequently updated with documents from 2021. We also downloaded a later version on March 09th, 2022, and merged both versions using each document’s doc_id field to produce the corpus used in this analysis. We divided documents by publication year and then preprocessed each using spacy’s en_core_web_sm model [26]. We replaced each tagged word or phrase with its corresponding entity type and entity id for every sentence that contained an annotation. Then, we used spacy to break sentences into individual tokens and normalized each token to its root form via lemmatization. After preprocessing, we used every sentence to train multiple natural language models designed to represent words based on their context.

#### Biomedical Preprints

BioRxiv [5] and MedRxiv [6] are repositories that contain preprints for the life science community. MedRxiv mainly focuses on preprints that mention patient research, while bioRxiv focuses on general biology. We downloaded a snapshot of both resources on March 4th, 2022, using their respective Amazon S3 bucket [27,28]. This snapshot contained 172,868 BioRxiv preprints and 37,517 MedRxiv preprints. These resources allow authors to post multiple versions of a single preprint. To prevent duplication bias, we filtered every preprint to its most recent version and sorted each preprint into its respective posted year. Unlike Pubtator Central, these filtered preprints do not contain any annotations. Therefore, we used TaggerOne [22] to tag every chemical and disease entity and GNormplus [23] to tag every gene and species entity for our preprint set. Once tagged, we used spacy to preprocess every preprint as described in our Pubtator Central section.

### Constructing Word Embeddings for Semantic Change Detection

Word2vec [29] is a natural language processing model designed to model words based on their respective neighbors in the form of dense vectors. This suite of models comes in two forms, a skipgram model and a continuous bags of words (CBOW) model. The skipgram model generates these vectors by having a shallow neural network predict a word’s neighbors given the word, while the CBOW model predicts the word given its neighbors. We used the CBOW model to construct word vectors for each year. Despite the power of these word2vec models, these models are known to differ both due to randomization within year and year-to-year variability across years [30,31,32,33]. To control for run-to-run variability, we examined both intra-year and inter-year relationships. Each year, we trained ten different CBOW models using the following parameters: vector size of 300, 10 epochs, minimum frequency cutoff of 5, and a window size of 16 for abstracts. Every model has its own unique vector space following training, making it diffcult to compare two models without a correction step. We used orthogonal Procrustes [34] to align models. We aligned all trained CBOW models for the Pubtator Central dataset to the first model trained in 2021. Likewise, we aligned all CBOW models for the BioRxiv/MedRxiv dataset to the first model trained in 2021. We used UMAP [35] to visually examine the aligned models. We trained this model using the following parameters: cosine distance metric, random_state of 100, 25 for n_neighbors, a minimum distance of 0.99, and 50 n_epochs.

### Detecting semantic changes across time

Once word2vec models are aligned, the next step is to detect semantic change.

Semantic change events are often detected through time series analysis [36]. We constructed a time series sequence for every token by calculating its distance within a given year (intra-year) and across each year (inter-year). We used the model pairs constructed from the same year to calculate an intra-year distance. Then, we calculated the cosine distance between each token and its corresponding counterpart for every generated pair. Cosine distance is a metric bounded between zero and two, where a score of zero means two vectors are the same, and a score of two means both vectors are different. For the inter-year distance, we used the Cartesian product of every model between two years and calculated the distance between tokens in the same way as the intra-year distance. Following both calculations, we combined both metrics by taking the ratio of the average inter-year distance over the average intra-year distance. Through this approach, tokens with high intra-year instability will be penalized and vice-verse for more stable tokens. Along with token distance calculations, it has been shown that including token frequency improves results compared to using distance alone [37]. We calculated token frequency as the ratio of token frequency in the more recent year over the frequency of the previous year. Then, we combined both the frequency and distance ratios to make the final metric.

Following time series construction, we performed change point detection, which is a process that uses statistical techniques to detect abnormalities within a given time series. We used the CUSUM algorithm [11] to detect these abnormalities. This algorithm uses a rolling sum of the differences between two timepoints and checks whether the sum is greater than a threshold. A changepoint is considered to have occurred if the sum is greater than a threshold. We used the 99th percentile on every generated timepoint as the threshold. Then, we ran the CUSUM algorithm using a drift of 0 and default settings for all other parameters.

## Results

### Models can be aligned and compared within and between years

We examined how the usage of tokens in biomedical text changes over time. Our evaluation was derived from machine learning models designed to predict the actual token given a portion of its surrounding tokens. Each token was represented as a vector in a coordinate space constructed by these models. However, training these models is stochastic, which results in arbitrary coordinate spaces. Model alignment is an essential step in allowing word2vec models to be compared [38,39]. Before alignment, each model has its own unique coordinate space (Figures 1A), and each word is represented within that space (Figure 1B). Alignment projects every model onto a shared coordinate space (Figure 1C), enabling direct token comparison. We randomly selected 100 tokens to confirm that alignment worked as expected. In aligned models, tokens in the global spcae were more similar to themselves within year than between years, while identical tokens in unaligned models were completely distinct (Figure 1D). Local distances were unaffected by alignment (Figure 1D), as token-neighbor distances were unaffected by the alignment procedure.

**Figure 1:**
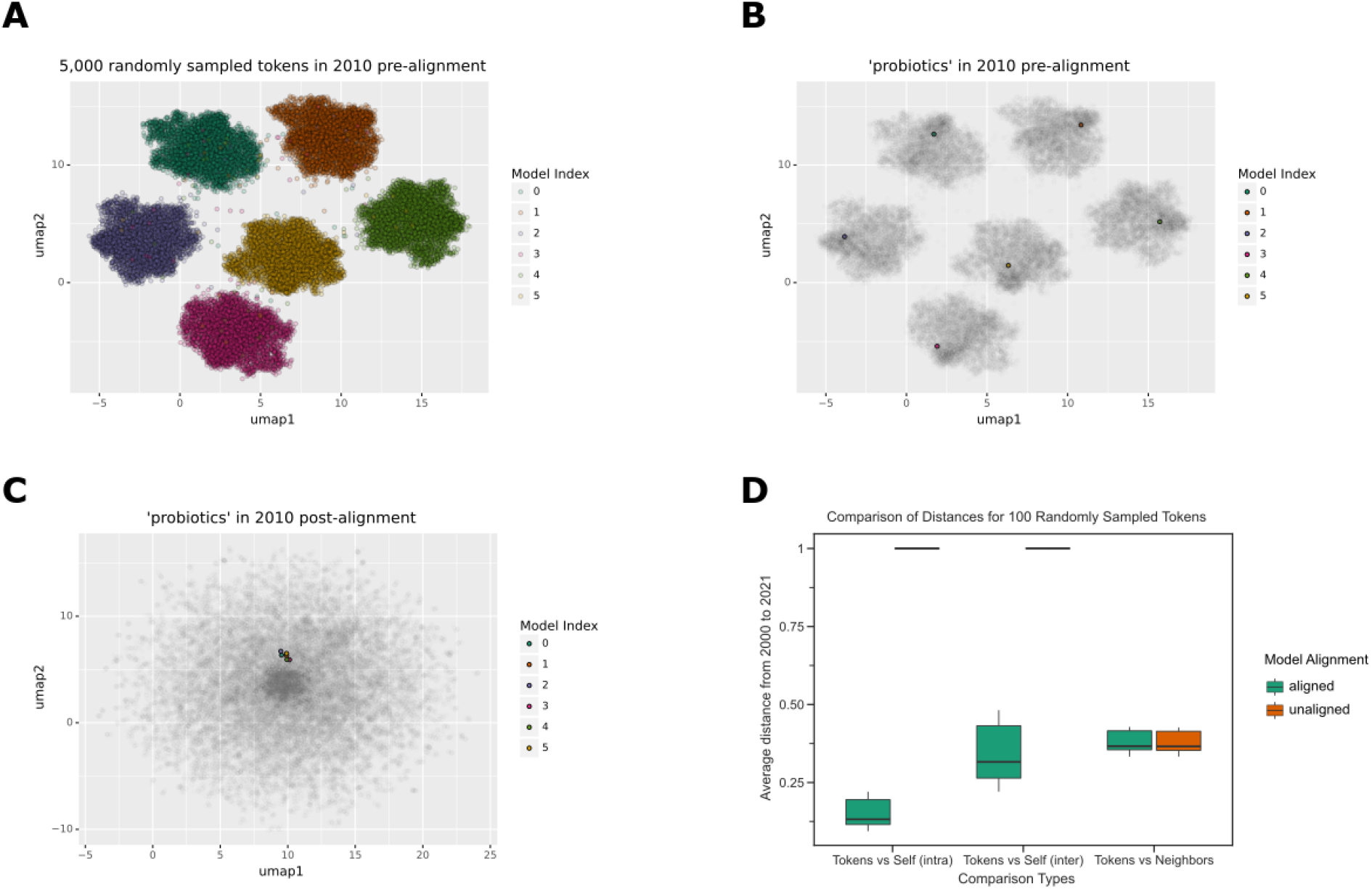
A. Without alignment, each word2vec model has its own coordinate space. This is a UMAP visualization of 5000 randomly sampled tokens from 5 distinct Word2Vec models trained on the text published in 2010. Each data point represents a token, and the color represents the respective Word2Vec model. B. The highlighted token ‘probiotics’ shows up in its respective clusters. Each data point represents a token, and the color represents the Word2Vec model. C. After the alignment step, the token ‘probiotic’ is closer in vector space. Each data point represents a token, and the color represents the different Word2Vec models. D. In the global coordinate space, token distances appear to be vastly different without alignment, but become closer upon alignment, while local distances, evaluated using neighbors, are unaffected. This boxplot shows the average distance of 100 randomly sampled tokens shared in every year from 2000 to 2021. The x-axis shows the various groups being compared (tokens against themselves via intra-year and inter-year distances and tokens against their corresponding neighbors. The y axis shows the averaged distance for every year.

The landscape of biomedical publishing has changed rapidly during the period of our dataset. The texts for our analysis were open access manuscripts available through PubMed Central. The growth in the amount of available text and the uneven adoption of open access publishing during the interval studied was expected to induce changes in the underlying machine learning models, making comparisons more diffcult. We found that the number of tokens available for model building, i.e., those in PMC OA, increased dramatically during this time (Figure 2A). This was expected to create a pattern where models trained in earlier years were more variable than those from later years simply due to the limited sample size in early years. We aimed to correct for this change in the underlying models by developing a statistic that, instead of using pairwise comparisons of token distances between individual models, integrated multiple models for each year by comparing tokens’ intra- and inter-year variabilities. We defined the statistic as the ratio of the average distance between two years over the sum of the average distance within each year respectively.

**Figure 2:**
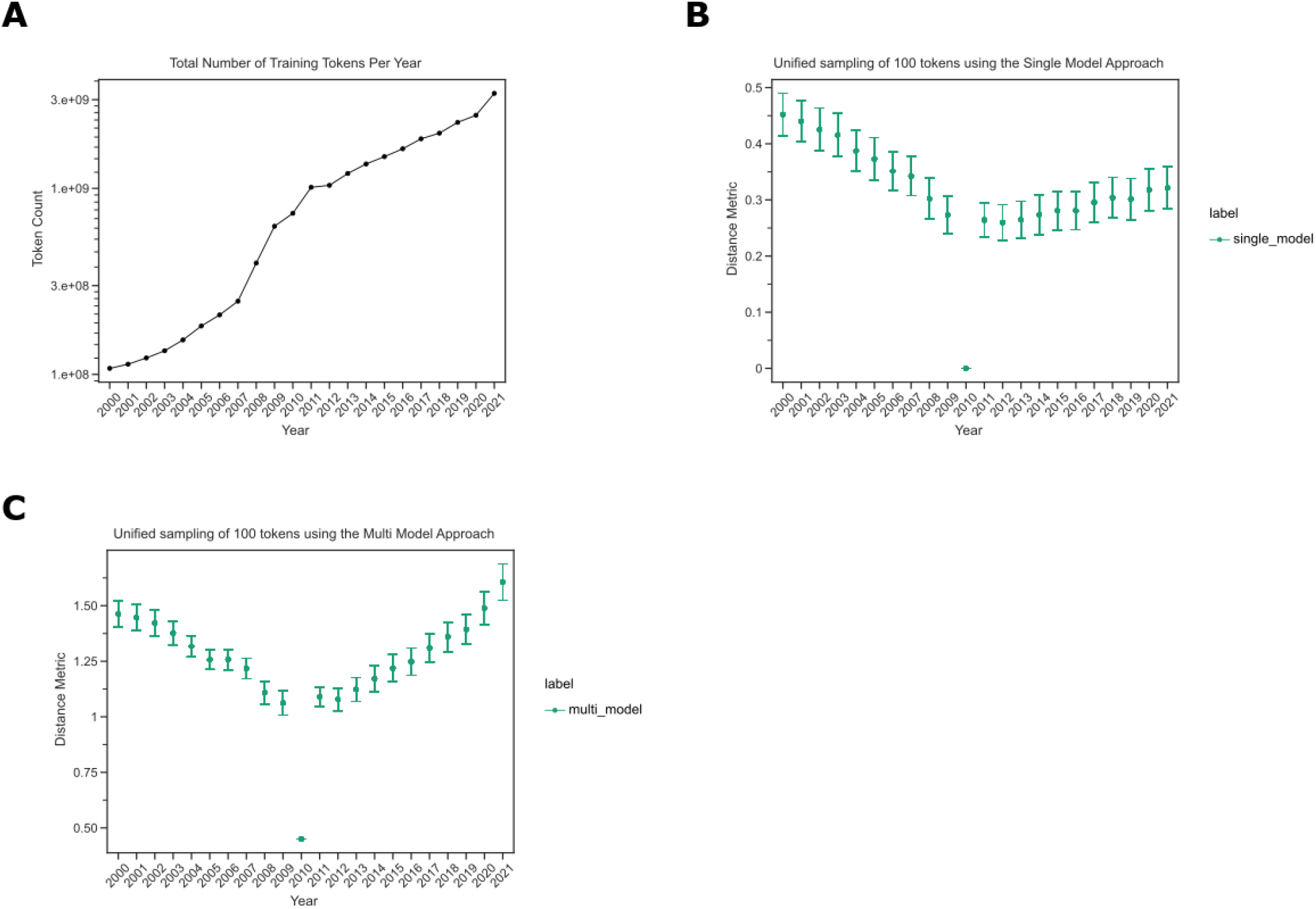
A. The number of tokens our models have trained on increases over time. This line plot shows the number of unique tokens seen by our various machine learning models. The x-axis depicts the year and the y-axis shows the token count. B. Earlier years compared to 2010 have greater distances than later years. This confidence interval plot shows the collective distances obtained by sampling 100 tokens that are present from every year using a single model approach. The x-axis shows a given year and the y-axis shows the distance metric. C. Later years have a lower intra-distance variability compared to the earlier years. This confidence interval plot shows the collective distances obtained by sampling 100 tokens that are present from every year using our multi-model approach. The x-axis shows a given year and the y-axis shows the distance metric.

We expected most tokens to undergo minor changes from year to year, while substantial changes likely suggested model drift as opposed to true linguistic change. We measured the extent to which tokens differed from themselves using the standard single-model approach and our integrated statistic. We filtered the token list to only contain tokens present in every year and compared their distance to the midpoint year, 2010, using the single-model and integrated-models strategies. We found that distances tended were markedly larger in the earliest years, where we expected models to be least stable, using the traditional approach (Figure 2B). The integrated model approach did not display the same pattern in the earliest years (Figure 2C). Both trends reinforce that training on smaller corpora will lead to high variation and that an integrated model strategy is needed [32]. Based on these results, we used the integrated-model strategy to calculate inter-year token distances for the remainder of this work.

### Terms exhibit detectable changes in usage

We next sought to identify tokens that changed during the 2000-2021 interval for the text from PubMed Central’s Open Access Corpus (PMCOA) and the 2015-2022 interval for our preprint corpus. We performed change point detection using the CUSUM algorithm with distances calculated with the integrated-model approach to correct for systematic differences in the underlying corpora. We found 41281 terms with a detected change point from PMCOA and 2266 terms from preprints (Figures 3A and 3B), and the vast majority (38019 for PMCOA and 2260 for preprints) had just a single change-point.

**Figure 3:**
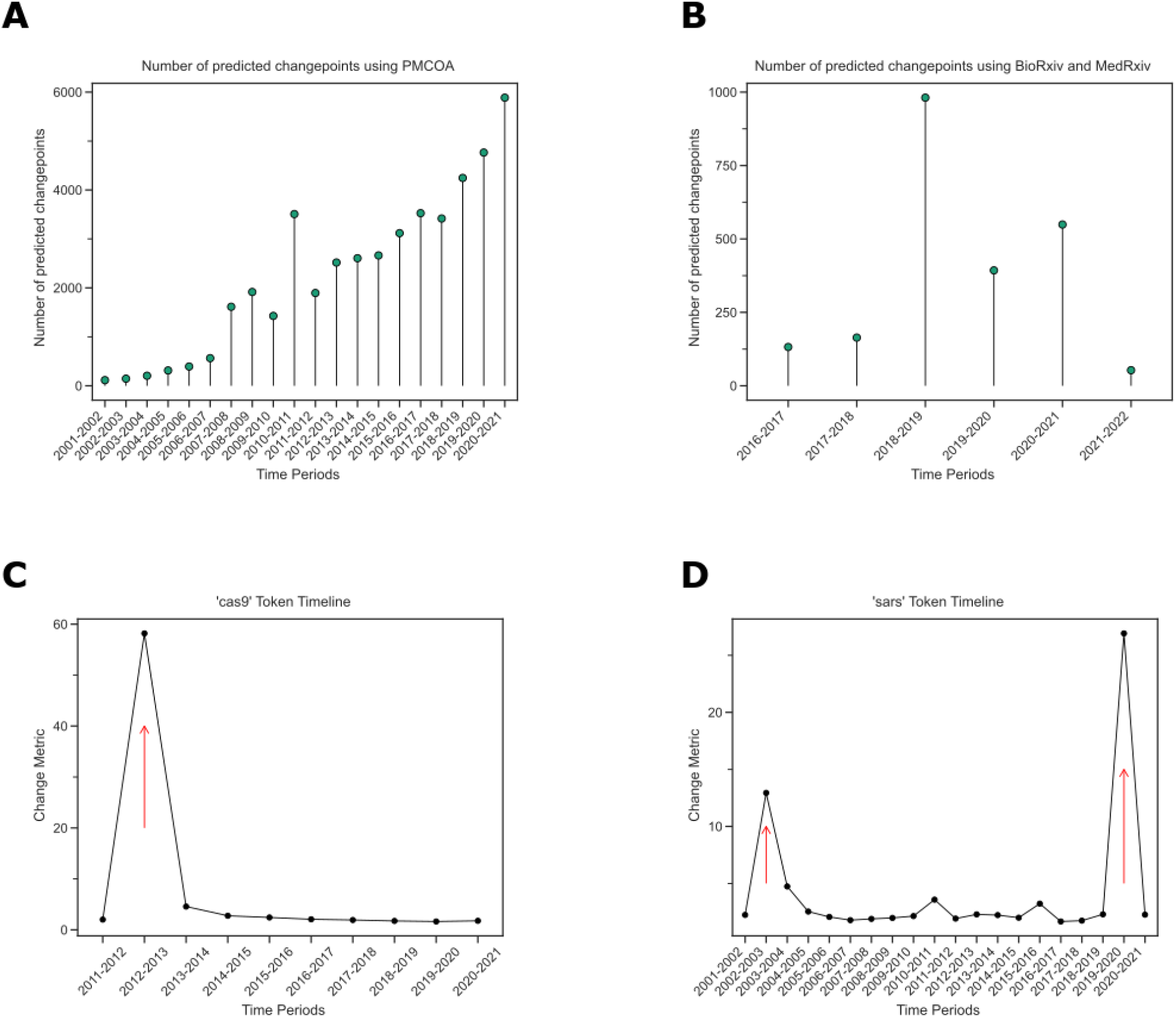
A. The number of change points increases over time in PMCOA. The x-axis shows the various time periods, while the y-axis depicts the number of detected change points. B. Regarding preprints, the greatest number of change points was during 2018-2019. The x-axis shows the various time periods, while the y-axis depicts the number of detected change points. C. The token ‘cas9’ was detected to have a change point at 2012-2013. The x-axis shows the time period since the first appearance of the token, and the y-axis shows the change metric. D. ‘sars’ has two detected change points within the PMCOA corpus. The x-axis shows the time period since the first appearance of the token, and the y-axis shows the change metric.

We explored individual change points. We detected one in PMCOA for ‘cas9’ from 2012 to 2013 (Figure 3C). Before the change point, its closest neighbors were related genetic elements (e.g., ‘cas’1-3). After the change point, its closest neighbors became terms related to targeting, sgRNA, and gRNA, as well as other genome editing strategies, ‘talen’ and ‘zfns’ (Table 1). For some terms, we detected multiple change points within the studied interval. We detected change points for ‘SARS’ from 2002 to 2003 and 2019 to 2020 (Figure 3D), consistent with the emergences of SARS-CoV [40] and SARS-CoV-2 [41,42] as observed human pathogens. We found miscellaneous neighbors before each change point, with use consistent with the acronym for Severe Acute Respiratory Syndrome after each (Tables 2 and 3).

**Table 1:**
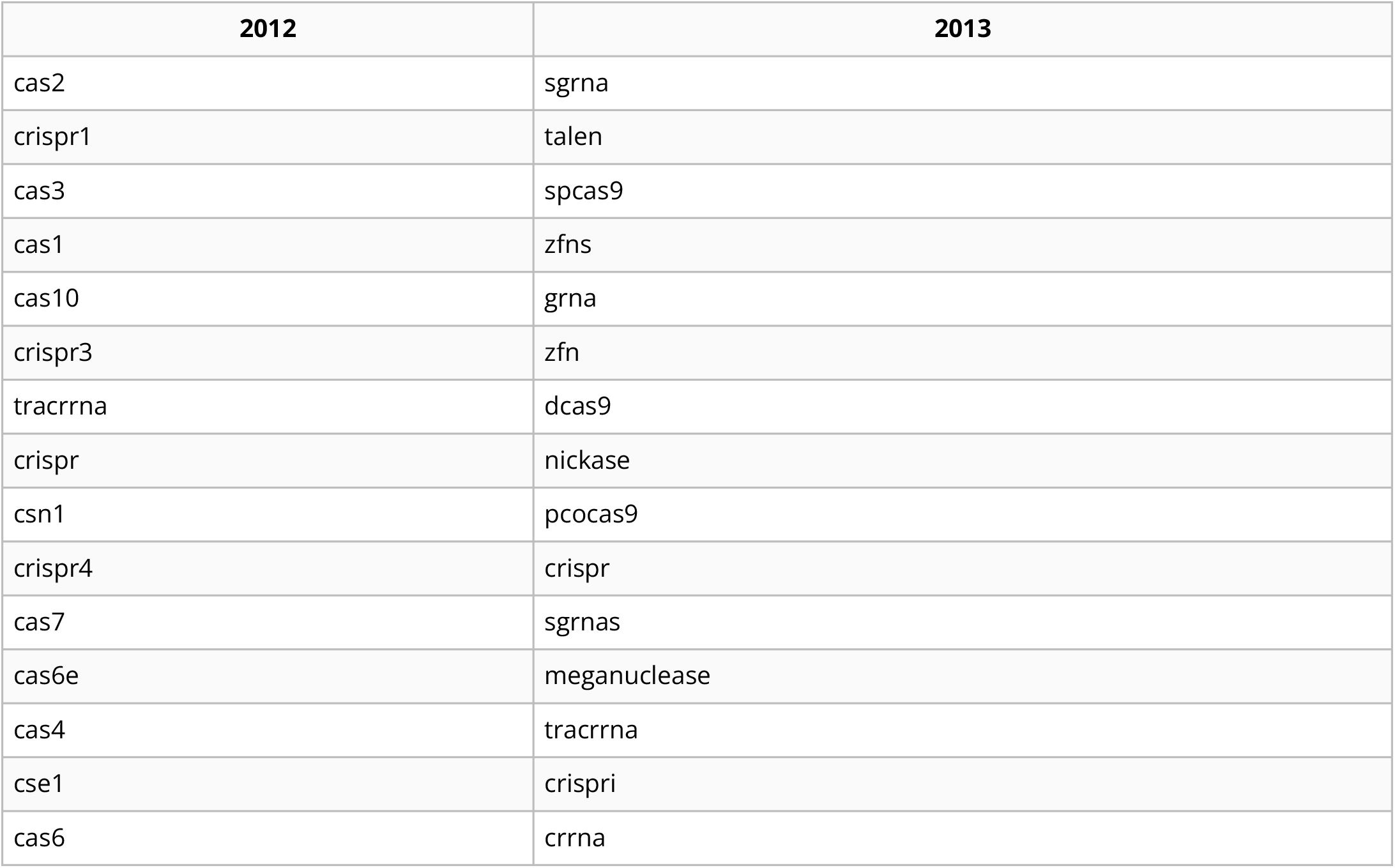
The fifteen most similar neighbors to the token ‘cas9’ for the years 2012 and 2013.

**Table 2:**
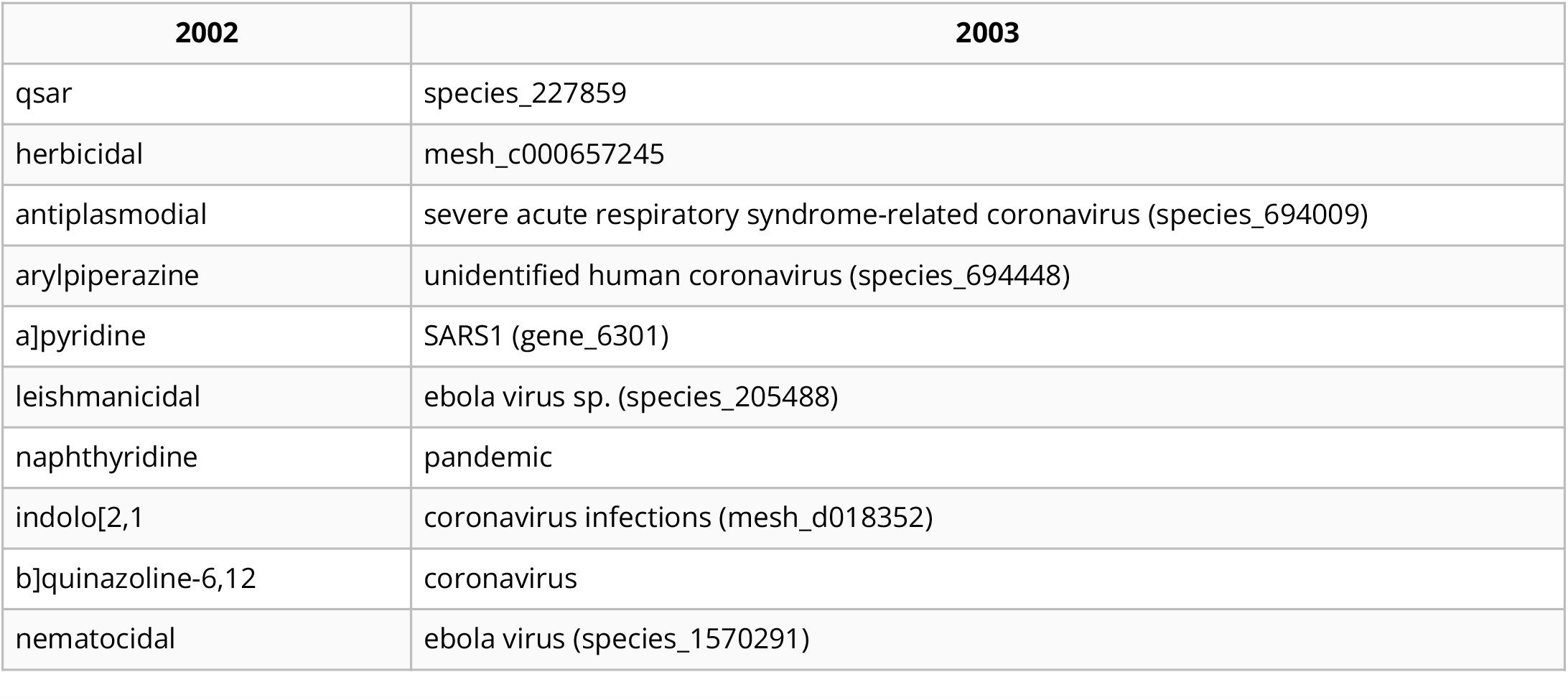
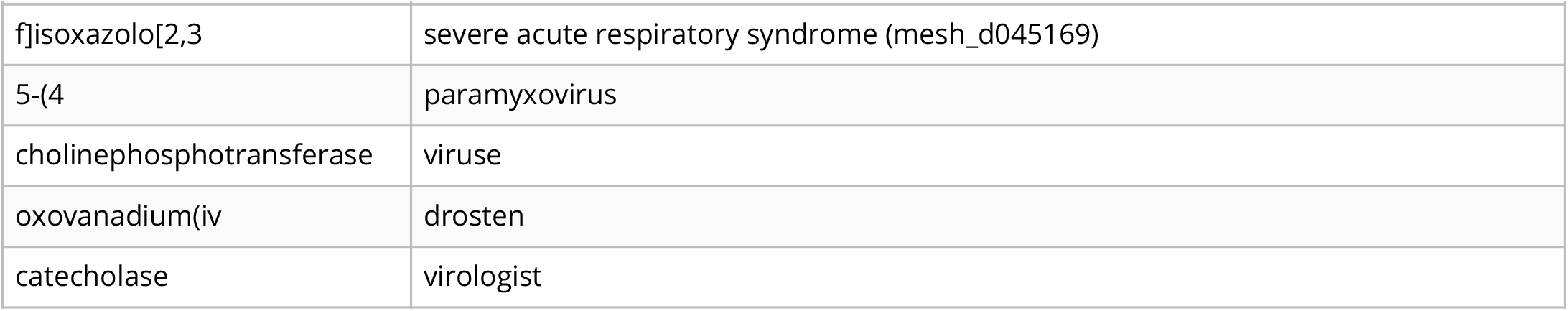
The fifteen most similar neighbors to the token ‘sars’ for the years 2002 and 2003.

**Table 3:**
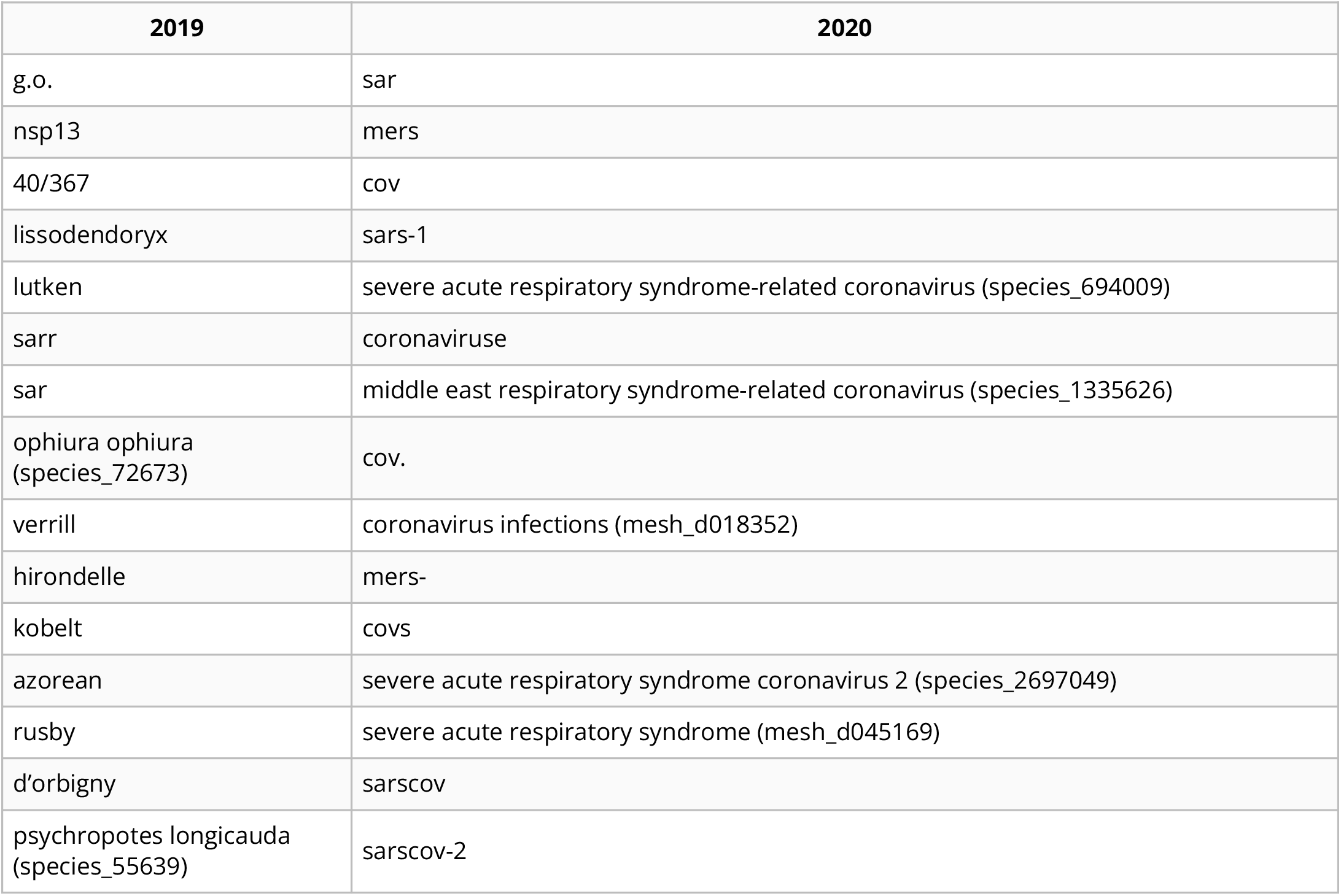
The fifteen most similar neighbors to the token ‘sars’ for the years 2002 and 2003.

Out of all change points, we observed 200 tokens with at least one change point in each corpus. Only 25 of the 200 terms were detected to have simultaneous changes between the preprint and PMCOA corpora. We examined the overlap of detected change points between preprints and published articles. Many of these 25 were related to the COVID-19 pandemic (Supplementary Table S1). The complete set of detected change points is available for further analysis (see Data Availability and Software).

### The word-lapse application is an online resource for manual examination of biomedical tokens

We constructed an online application that allows users to examine how tokens change through time. The application supports token input as text strings or as MeSH IDs, Entrez Gene IDs, and Taxonomy IDs. Users might elect to explore the term ‘pandemic’, for which we detected a change point between 2019 and 2020. Users can examine the token’s nearest neighbors through time (Figure 4A). Using the token ‘pandemic’ as an example, users can observe that ‘epidemic’ remains similar through time, but taxid:114727 (the H1N1 subtype of influenza) only entered the nearest neighbors with the swine flu pandemic in 2009 and that MeSH:C000657245 (COVID-19) appears in 2020. The application also shows a frequency chart depicting how often the particular token is used each year (Figure 4B), which can be displayed as a raw count or adjusted by the total size of the corpus. When change points are detected, they are indicated on this panel (Figure 4B). The final visualization shows the union of the nearest 25 neighbors from each year ordered by the number of years that neighbor was present (Figure 4C). This visualization has a comparison function that allows users to examine differences between years. All functionalities are fully supported across the PMCOA and preprint corpora, and users can toggle between the two.

**Figure 4:**
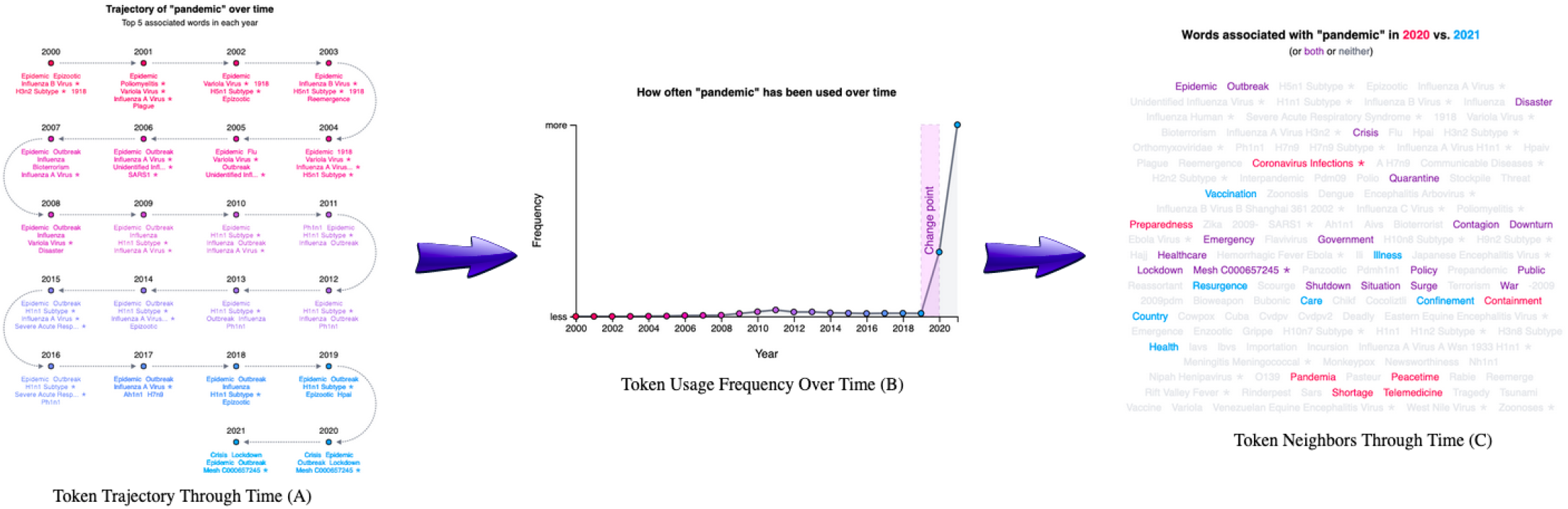
A. The trajectory visualization of the token ‘pandemic’ through time. It starts at the first mention of the token and progresses through each subsequent year. Every data point shows the top five neighbors for the respective token. B. The usage frequency of the token ‘pandemic’ through time. The x-axis shows the year, and the y-axis shows the frequency for each token. C. A word cloud visualization for the top 25 neighbors for the token ‘pandemic’ each year. This visualization highlights each neighbor from a particular year and allows for the comparison between two years. Tokens in purple are shared within both years, while tokens in red or blue are unique to their respective year.

## Discussion and Conclusion

Language is rapidly evolving, and the usage of words changes over time with words assimilating new meanings or associations [1]. Some efforts have been made to study semantic change using biomedical text [18,19,20]; however, no such work has examined the changes evident in both pre-publication peer-reviewed and preprinted biomedical text. We examined semantic change in both open-access biomedical corpora from Pubmed (PMCOA) and bioRxiv/MedRxiv for the 2000-2021 interval. Studies like this one have only become feasible recently with the rapid uptake of open access publishing. To be able to span two decades with very different data availability, we developed a novel statistic that incorporated multiple models using both inter- and intra-year distances. Without the correction, comparing between stable and unstable models is challenging as previously reported [32,43]. Our analysis revealed more than 41,000 different change points, including tokens such as ‘cas9’, ‘pandemic’, and ‘sars’. Many change points overlapping between PMCOA and preprints were related to COVID-19, indicating that the COVID-19 pandemic has been strong and immediate enough to induce rapid semantic change across both publishing paradigms.

As the amount of preprinted text grows, future work may be able to determine the consistency and time-lag of semantic change between preprint and pre-publication peer-reviewed text - potentially predicting future change in pre-publication peer-reviewed text. We developed a web application to enable users to investigate individual tokens - automatic approaches that use orthogonal metrics to estimate change point validity would make analysis at scale more straightforward. Furthermore, including corpora such as the arXiv [44] or psyArXiv [45] repositories may reveal consistencies across a broader swath of fields or within-field analyses may reveal the earliest starting points of semantic changes that ultimately sweep through biomedicine.

## Data Availability and Software

An online version of this manuscript is available under a Creative Commons Attribution License at https://greenelab.github.io/word_lapse_manuscript/. The source for the research portions of this project is licensed under the BSD-2-Clause Plus Patent at https://github.com/greenelab/biovectors.

Our Word Lapse website can be found at https://greenelab.github.io/word-lapse, and the code for the website is available under a BSD-3 Clause at https://github.com/greenelab/word-lapse. Full-text access for the bioRxiv repository is available at https://www.biorxiv.org/tdm. Full-text access for the medRxiv repository is available at https://www.medrxiv.org/tdm. Access to Pubtator Central’s Open Access subset is available on NCBI’s FTP server at https://ftp.ncbi.nlm.nih.gov/pub/lu/PubTatorCentral/.

## Acknowledgments

This work was supported by the Gordon and Betty Moore Foundation under award GBMF4552 and the National Institutes of Health’s National Human Genome Research Institute under award R01 HG010067 to CSG. The funders had no role in the study design, data collection, and analysis, decision to publish, or preparation of the manuscript.

## Competing Interest

DNN performed this work as a Ph.D. student at the University of Pennsylvania. He is now employed by Digital Science. The authors report no other competing interests for this paper.

## Supplemental Tables

**Table S1:** The intersection of changepoints found between published papers and preprints.

| Token | Changepoint |
| --- | --- |
| lockdown | 2019-2020 |
| 2021 | 2020-2021 |
| distancing | 2019-2020 |
| 2019 | 2018-2019 |
| ace2 | 2019-2020 |
| pandemic | 2019-2020 |
| 2020 | 2019-2020 |
| coronavirus | 2019-2020 |
| bcl2a1 | 2018-2019 |
| peak3 | 2020-2021 |
| 3.6.2 | 2019-2020 |
| quarantine | 2019-2020 |
| cobl | 2020-2021 |
| injectrode | 2020-2021 |
| nrc3 | 2020-2021 |
| 4.0.5 | 2020-2021 |
| TMPRSS2 (gene_7113) | 2019-2020 |
| n262 | 2019-2020 |
| bin1 | 2017-2018 |
| n3c | 2020-2021 |
| tip1 | 2020-2021 |
| omicron | 2020-2021 |
| pangolin | 2019-2020 |
| adrn | 2020-2021 |
| seir | 2019-2020 |

## Notes

https://github.com/greenelab/word_lapse_manuscript

https://github.com/greenelab/word-lapse

https://github.com/greenelab/biovectors

### Competing Interest Statement

The authors have declared no competing interest.

